# Hypoxic volatile metabolic markers in the MDA-MB-231 breast cancer cell line

**DOI:** 10.1101/2023.03.02.530779

**Authors:** Theo Issitt, Matthew Reilly, Sean T. Sweeney, William J. Brackenbury, Kelly Redeker

## Abstract

Hypoxia in disease describes persistent low oxygen conditions, observed in a range of pathologies, including cancer. In the discovery of biomarkers in biological models, pathophysiological traits present a source of translatable metabolic products for the diagnosis of disease in humans. Part of the metabolome is represented by its volatile, gaseous fraction; the volatilome. Human volatile profiles, such as those found in breath, are able to diagnose disease, however accurate volatile biomarker discovery is required to target reliable biomarkers to develop new diagnostic tools. Using custom chambers to control oxygen levels and facilitate headspace sampling, the MDA-MB-231 breast cancer cell line was exposed to hypoxia (1% oxygen) for 24 hours. The maintenance of hypoxic conditions in the system was successfully validated over this time period. Targeting and non-targeting gas chromatography mass spectrometry approaches revealed four significantly altered volatile organic compounds when compared to control cells. Three compounds were actively consumed by cells: methyl chloride, acetone and n-Hexane. Cells under hypoxia also produced significant amounts of styrene. This work presents a novel methodology for identification of volatile metabolisms under controlled gas conditions with novel observations of volatile metabolisms by breast cancer cells.

## Introduction

The human ‘volatilome’ describes the production and metabolism by the human body of small, carbon-containing compounds called Volatile Organic Compounds (VOCs) which are gaseous at room temperature and pressure (1, 2). VOCs can be found in abundance in the breath and are reflective of processes within the body (2, 3). Although fluctuations of VOCs vary between individuals and throughout the day, disease specific ‘volatile fluxes’, or biomarkers, could provide opportunities to non-invasively diagnose disease, monitor treatment and measure bodily functions (3, 4).

The clinical potential of VOCs in diagnosis has been shown by a number of published breath studies (3). Diagnostic accuracy using breath VOC biomarkers has been achieved for a wide range of conditions, including various types of cancer (3, 5), liver disease (6), diabetes (7), transplant rejection (8), infections of the lung (3, 9), liver function (using labelled VOCs) (10) and other conditions (3). Each study may independently achieve high sensitivity of disease detection (i.e. > 90%) but the reported compounds often do not translate between studies, slowing clinical application through conflicting and confounding results (3). However, our recent meta-analysis has shown underlying trends in chemical functional groups from published studies supporting potential clinical application (3). It is clear that in order to identify effective biomarkers more targeted methodological approaches are required to overcome variability (3, 11).

VOC profiles from cell types associated with pathological conditions have been identified, for example, differences between breast (4, 12), liver (13) and mesothelioma (14) cancer cell lines. However, cellular VOC studies tend to be non-stressed cells in high (21%, atmospheric) oxygen conditions, which is not consistent with many disease or normal physiological states. To accelerate biomarker discovery, we propose models of pathophysiological stress. For example; stress from reactive oxygen species (ROS) induces alkane release in breast cancer cells (15), VOCs which have been observed in the breath of ROS associated conditions (3).

Hypoxia is a persistent reduction in oxygen from normal physiological conditions (normoxia). It is characteristic of a range of diseases, including, pulmonary hypertension (16) and cancer (17). It induces a range of metabolic alterations, including reduction in adenosine triphosphate generation and inhibition of fatty-acid desaturation through hypoxia inducible factor activity (16, 17, 18), which can produce alterations in a range of associated breath volatiles (19, 20). Despite its relevance to pathophysiology, hypoxic volatiles have yet to be investigated in vitro. This is partially due to the challenges associated with development of a headspace sampling tool which can maintain an hypoxic environment. While volatile compounds in the available, limited, published studies associated with hypoxia show variation in breath (19, 20), translatable studies are required for target biomarker discovery.

Biomarker discovery in appropriate biological models can accelerate clinical delivery by identifying and allowing targeted analytical approaches, separating methodical challenges from pathology, and improving sensitivity. Multi-timepoint sampling and approaches considering local environment will also accelerate clinical application of breath diagnostics and consideration of methodological challenges around clinical application should drive experimental design. We have previously demonstrated a platform and method for both identification of VOC metabolisms in cellular headspace over time and VOC changes in response to cellular stress (4). However, models of pathological conditions require further investigation to ensure biomarker discovery is translatable from cell to human.

One of the primary sources of variance within the published literature revolves around methodology. Methods of breath VOC analysis can be split into 3 main sections where variability between studies can arise: initial collection, sample transfer and analytical approach. There are many effective breath collection methods for analysis of VOCs, such as simply breathing into a specialised bag or use of specialised technologies (11, 21). Many studies use single time point collection (3), considering presence verses absence, which can miss valuable metabolic information, particularly volatile uptake, driven via chemical reactions reflective of cellular state or through cellular metabolism. Furthermore, variability in local environment influences and reduces reported outcome precision (3, 21, 22) and approaches should consider sampling the environment (i.e. - ambient air) along with breath (11). A sample, once collected, is then transferred, either directly or indirectly (such as through chemical traps) to an analytical instrument. There are two main analytical approaches for discovery and accurate detection of VOCs: targeted and non-targeted. Non-targeted approaches, investigating the breath of patients, are capable of identifying relatively concentrated material (ppbv) whereas targeted approaches generally are capable of quantifying lower concentrations (pptv). Non targeted approaches therefore may miss changes in important, low-concentration compounds, while targeted approaches can only look only for a limited number of known compounds of interest, reducing discovery potential.

Here, we apply hypoxic stress to well-studied breast cancer cell lines with the intent of identifying process and disease-linked physiological volatile metabolisms specifically linked to low oxygen conditions, so that more accurate diagnostic tools can be developed and applied in the clinic. We utilised both targeted and non-targeted assays after sampling with a static headspace method that accounts for the ambient air background and allows quantification of cellular uptake of VOCs. We predicted that upon successful maintenance of a hypoxic environment, cellular VOC profiles from hypoxic versus hyperoxic cellular models would alter significantly.

## Methods

Methods for culture of MDA-MB-231 cells, headspace sampling from custom chambers and GC-MS analysis have been previously described in detail (4).

### Cell culture

MDA-MB-231 breast cancer cells (a gift from Professor Mustafa Djamgoz, Imperial College London) were grown in Dulbecco’s Modified Eagle Medium (DMEM, Thermo Scientific, Waltham, MA, USA), 25 mM glucose, supplemented with L-glutamine (4 mM) and 5% foetal bovine serum (Thermo Scientific, Waltham, MA, USA). Cell culture medium was supplemented with 0.1 mM NaI and 1 mM NaBr (to model physiological availability of iodine and bromide). All cells were grown at 37 °C with 5% CO_2_.

Prior to volatile collection, cells were trypsinised, and 500,000 cells were seeded into 8 mL complete media in 10 cm polystyrene cell culture dishes. Cells were then allowed to attach for 3-4 h, washed with warm PBS and 6 mL treatment media was applied. Volatile headspace sampling was performed 24 h later.

### Induction of the hypoxic environment and VOC headspace sampling

Cells were placed in static headspace chambers as previously described [4] with new, clean silicon gaskets. Low oxygen, hypoxic gas (1% O_2_, 5% CO_2_, 94% N_2_; purchased from BOC Specialty Gases, Woking, UK) was flushed through the chambers at a rate of 4L/min for 10 min (chamber volume = 25L). Chambers were then closed and placed at 37 °C for 2 hours to allow residual oxygen in the media to equilibrate with chamber headspace. Chambers were then flushed again at a rate of 4L/min for 10 min, sealed and returned to 37 °C.

After a further 24 hours, chambers were flushed again at a rate of 4L/min for 10 min. 15 ml of gas standards (MeCl, 520 ppb (parts per billion); MeBr, 22 ppb; MeI, 26 ppb; DMS, 110 ppb; CFC-11, 400 ppb and CH_3_Cl_3_, 110ppb; BOC Specialty Gases, Woking, UK) were then injected into the chambers through a butyl seal and time zero sample taken. Injected compounds are either known metabolites for cancer cells, or internal standards (CFC-11) for the analysis and quantification of metabolism. Final chamber concentrations were similar to environmental concentrations, e.g MeCl, 1.2 ppb and MeBr 0.05 ppb, particularly more polluted urban spaces (23). Injected gases are the same as those used for calibration. Compounds not injected but detected at first time point, due to residual presence from laboratory air, (including isoprene, acetone, 2-MP, 3-MP and n-hexane) were quantified. Two time zero (T0) samples were taken using an evacuated 500 mL electropolished stainless steel canister (LabCommerce, San Jose, USA) through fine mesh Ascarite^®^ traps (24), after which the chamber was resealed and left on a platform rocker on its slowest setting for 120 min, at which point two further air samples (T1) were collected. Duplicate samples were taken so that two analytical approaches could be performed (targeted and non-targeted MS).

Cells were removed from the chamber, washed with PBS twice and lysed in 500 µL RIPA buffer (NaCl (5 M), 5 mL Tris-HCl (1 M, pH 8.0), 1 mL Nonidet P-40, 5 mL sodium deoxycholate (10 %), 1 mL SDS (10%)) with protease inhibitor (Sigma-Aldrich, Roche; Mannheim, Germany). Protein concentration of lysates were determined using BCA assay (Thermo Scientific, Waltham, MA, USA).

Hypoxic condition media alone (1% O_2_) was treated exactly as described as above and no significant differences were detected (as confirmed by ANOVA) compared to media under normal lab air (21% O_2_). These media blank outcomes were combined and subtracted from cellular samples. Comparative controls include lab air blanks and those data available from the dataset and collection method published previously which created and quantified metabolic fluxes of volatile compounds from MDA-MB-231 under hyperoxic (lab air) conditions (4).

### GC-FID/TCD sampling and analysis (O2)

10 ml headspace samples were taken from chambers using an airtight syringe (10 ml, SGE, Trajan, Milton Keynes, UK). 1% O_2_ (BOC Specialty Gases, Woking, UK) was flushed through sealed chambers containing 6 ml DMEM as described for cell treatments. Samples were taken at 5 and then 10 min post initial flush. In order to replicate cell treatments, the chamber was then closed for 2 hours, then flushed for 10 min, after which an air sample was taken. A further 20 min flush with 1% O_2_ air was employed and the chamber was closed, placed at 37°C, and left to incubate for 24 hours, at which time the final sample was taken.

Air samples were immediately injected into an SRI 8610C Gas Chromatograph (SRI Instruments Europe GmbH, Torrance, CA, USA) and subsample of 0.2 ml was taken for analytical separation. Sample separation consisted sequentially of a Restek© PORAPAK Q porous polymer column (1.83 m length, 2.1 mm ID, 3.175 diameter thickness), a solenoid switching valve (for backflushing CO_2_) and a Restek© MOLECULAR 5A sieve column (0.91 m length, 2.1 mm ID, 3.175 diameter thickness). A flow of helium at 18 psi was used as a sample carrier gas. Flow of gas and column temperatures (50 °C) were maintained during separation. The valve was switched at 1.5 min to backflush the PORAPAK Q column. Measurement of compounds eluted from the MOLECULAR 5A sieve was achieved by using an SRI 8690-0030 Helium Ionisation Detector. SRI PeakSimple (version 453) software was used to generate a digital chromatograph for each sample and O_2_ was quantified by comparing the peak area to known standards.

The standard curve was developed by flushing 120ml Wheaton vials with butyl stoppers with pure nitrogen (BOC Gases, Woking, UK) for 30mins. 10ml of nitrogen only was injected to establish a background control. Because atmospheric air at sea level contains 21% O_2_, lab air was injected at 1%, 2%, 10%, 20% and 30% within the N_2_-filled vial to generate a standard curve consisting of 0%, 0.21%, 0.42%, 2.1%, 4.2% and 6.3% and 21% (lab air only). Peak areas were integrated using Graphpad (Prism), and Padé (1,1). Linear regression demonstrated an R squared value of 0.96.

### GC-MS analysis (Volatile Organic Compounds, VOCs)

Collected canister samples were transferred to a liquid nitrogen trap through pressure differential. Pressure change between beginning and end of “injection” was measured, allowing calculation of the moles of canister collected air injected. Sample in the trap was then transferred, via heated helium flow, to a Restek© (Bellefonte, PN, USA) PoraBond Q column (25 m length, 0.32 mm ID, 0.5-µm diameter thickness) connected to a quadrupole mass spectrometer (Aglient/HP 5972 MSD, Santa Clara, CA, USA). All samples were analysed with both targeted, select ion mode (SIM), and non-targeting (SCAN) mode, which quantified all ions between 45 and 200 amu. For details on SIM and significantly altered, identified SCAN compounds, see table 1. All samples were analyzed within 6 days of collection. The oven program for both SIM and SCAN analyses were identical and are as follows: 35 °C for 2 min, 10 °C/min to 115 °C, 1 °C/min to 131 °C and 25 °C min to 250 °C with a 5 min 30 sec hold. The quadrupole, ion source and transfer line temperatures were 280, 280 and 250 °C, respectively.

**Table 1.**
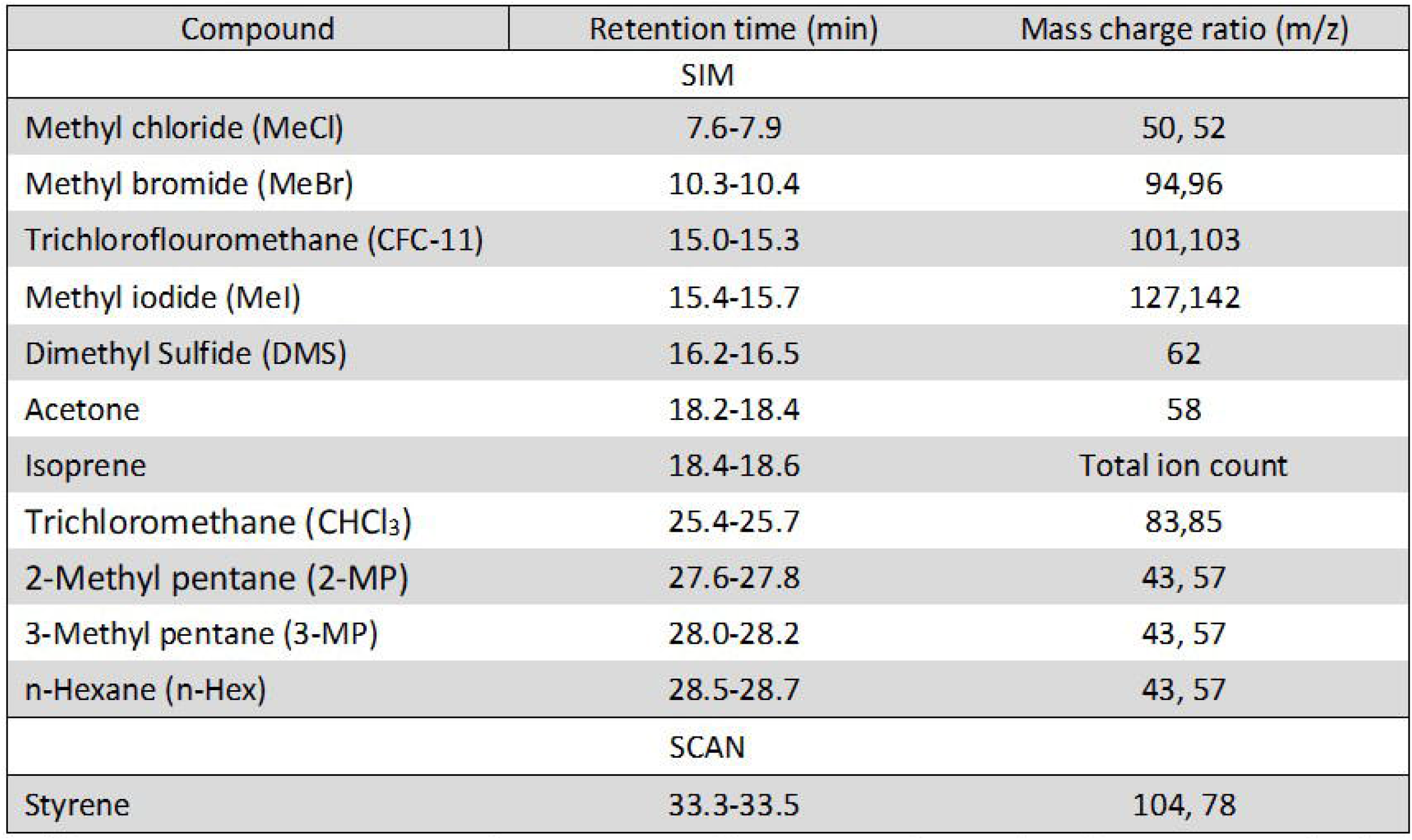
Retention times, mass charge ratios and GC-MS modes used to characterise individual VOCs. SIM and SCAN refer to select ion monitor and full scan (targeted and non-targeted) GC MS modes.

Calibration was performed using standard gases (BOC Specialty Gases, Woking, UK). Linear regression of calibration curves confirmed strong, positive linear relationships between observed compound peak areas and moles of gas injected for each VOC (r^2^ > 0.9 in all cases). For compounds not purchased in gaseous state (BOC Specialty gases, as above), 1–2 mL of compound in liquid phase was injected neat into butyl sealed Wheaton-style glass vials (100 mL) and allowed to equilibrate for 1 h. 1 mL of headspace air was then removed from neat vial headspace using a gas tight syringe (Trajan, SGE) and injected into the headspace of a second 100 mL butyl sealed Wheaton-style glass vial. This was then repeated, and 1 mL of the 2nd serial dilution vial was injected into the GCMS system with 29 mL of lab air to give ppb concentrations. This was performed for methanethiol (MeSH (SPEXorganics, St Neots, UK)), isoprene (Alfa Aesar, Ward Hill, MA, USA), acetone (Sigma-Aldrich, Burlington, MA, USA), 2- & 3-methyl pentane and n-hexane (Thermo Scientific, Waltham, MA, USA). Reported compounds detected by the GC-MS were confirmed by matching retention times and mass–charge (*m/z*) ratios with known standards.

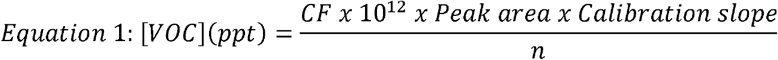

Equation 1 outlines the approach to calculating VOC concentrations in parts-per-trillion-by-volume, or pptv. Here Peak area refers to the combined peak areas for the mass-charge ratios identified in Table 1. Multiplying Peak areas by their associated calibration curves (*Calibration Slope*) generate molar amounts which, when divided by the number of moles of headspace air injected (*n*), generate a unitless (moles compound/moles of air) ratio. Pptv concentrations are then obtained by multiplying this unitless ratio by 1×10^12^. For clarity, part-per-billion-by-volume values would be obtained by multiplying the unitless ratios by 1×10^9^, or one billion. Sample VOC concentrations were then normalised to CFC-11 concentrations (240 parts-per-trillion-by-volume (pptv)) through multiplication by a “correction factor”, or *CF*, Equation 1). CFC-11 was used as an internal standard, since atmospheric concentrations of CFC-11 are globally consistent and stable (23). Quantification of Styrene was done as above but normalisation to CFC-11 was not possible under flushed, hypoxic conditions.

To account for differences in rates of cellular proliferation over 24hours, cellular results from GC-MS analyses were normalised to protein content at time of sampling using a BCA assay. When comparing media blanks to cellular assays results are reported in grams compound per petri dish per hour.

### Hydrogen peroxide (Amplex red) assay

Experiments were performed in phenol red free DMEM. DMEM containing 50 μM Amplex Red reagent (Thermo Scientific, Waltham, MA, USA) and 0.1 U/ml horse radish peroxidase (HRP, Thermo Scientific, Waltham, MA, USA) was added to cells in 12 well dishes (500 μl per well) for 15 min following 24 hours in hypoxic or control conditions. Fluorescence at 590 nm was measured with a plate reader (Clariostar, BMG, Ortenberg, Germany) and compared against a H_2_O_2_ standard curve for quantification.

## Results

### Chambers maintain low oxygen conditions over 24 hours

To confirm chambers maintained hypoxic conditions over 24 hours we sampled gas from chambers throughout our method, measuring O_2_. When flushed with reduced oxygen air (1%) for 5 minutes, oxygen levels rapidly fell from atmospheric 21% to between 6% and 2% (figure 1). After 10 min of reduced oxygen flushing, each chamber held less than 5%. Chambers left for 2 hours (120 mins) to allow media to equilibrate and flushed for 10 min revealed average O_2_ levels of 1.15% ± 1.03 (Ch 1), 1.34% ± 0.93 (Ch 2) and 1.98% ± 4.07 (Ch 3) respectively. Sealed chambers maintained low oxygen levels over 24 hours with average O_2_ levels of 1.31% ± 1.31 (Ch 1), 1.76% ± 1.02 (Ch 2) and 1.96% ± 0.28 (Ch 3) respectively.

**Figure 1.**
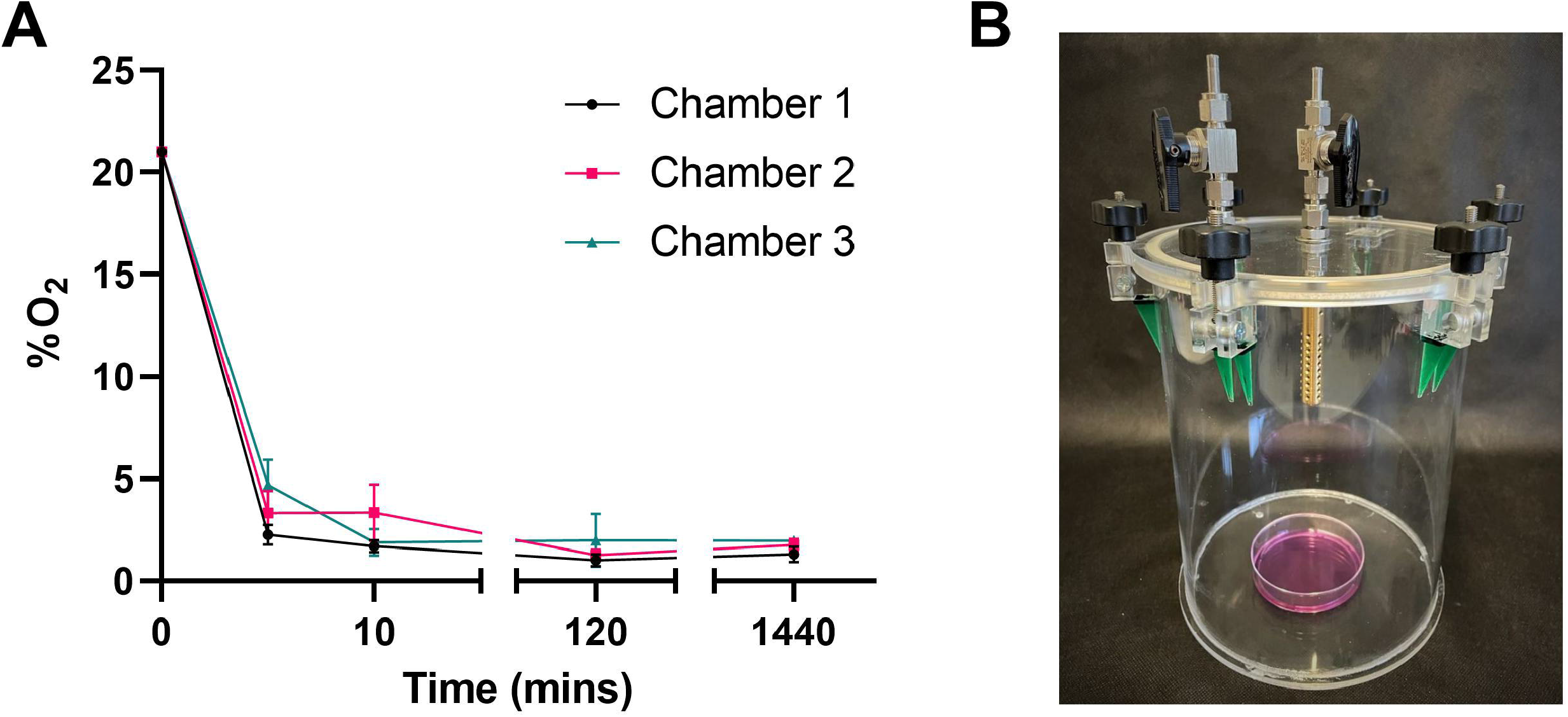
Chambers maintain hypoxic conditions over 24 hours. **(A)** Oxygen (O_2_) content in 3 custom made chambers containing 6ml media was measured following a 10 min flush, 2 hour dwell and another 10 min flush (20mins) with 1% O_2_, 5% CO_2_ gas mix. O_2_% was then measured following chambers being sealed for 1440 mins (24 hours). Mean ± SEM; n = 3. **(B)** Image of collection chamber.

### Hypoxia induces differing volatile fluxes in breast cancer cell line MDA-MB-231

Persistent hypoxia over 24 hours induced significant changes in flux for 3 targeted compounds (SIM analysis); MeCl, acetone and n-hexane (but not hexane isomers; 2-methyl pentane, or 3-methyl pentane), when compared to control (Figure 2A-C). MeCl was taken up by cells under hypoxia and released by cells under hyperoxic cell culture conditions. Acetone was not significantly produced or utilised by control cells but cells under hypoxia significantly consumed acetone. N-hexane was produced by hyperoxic control cells while those under hypoxia consumed hexane.

**Figure 2.**
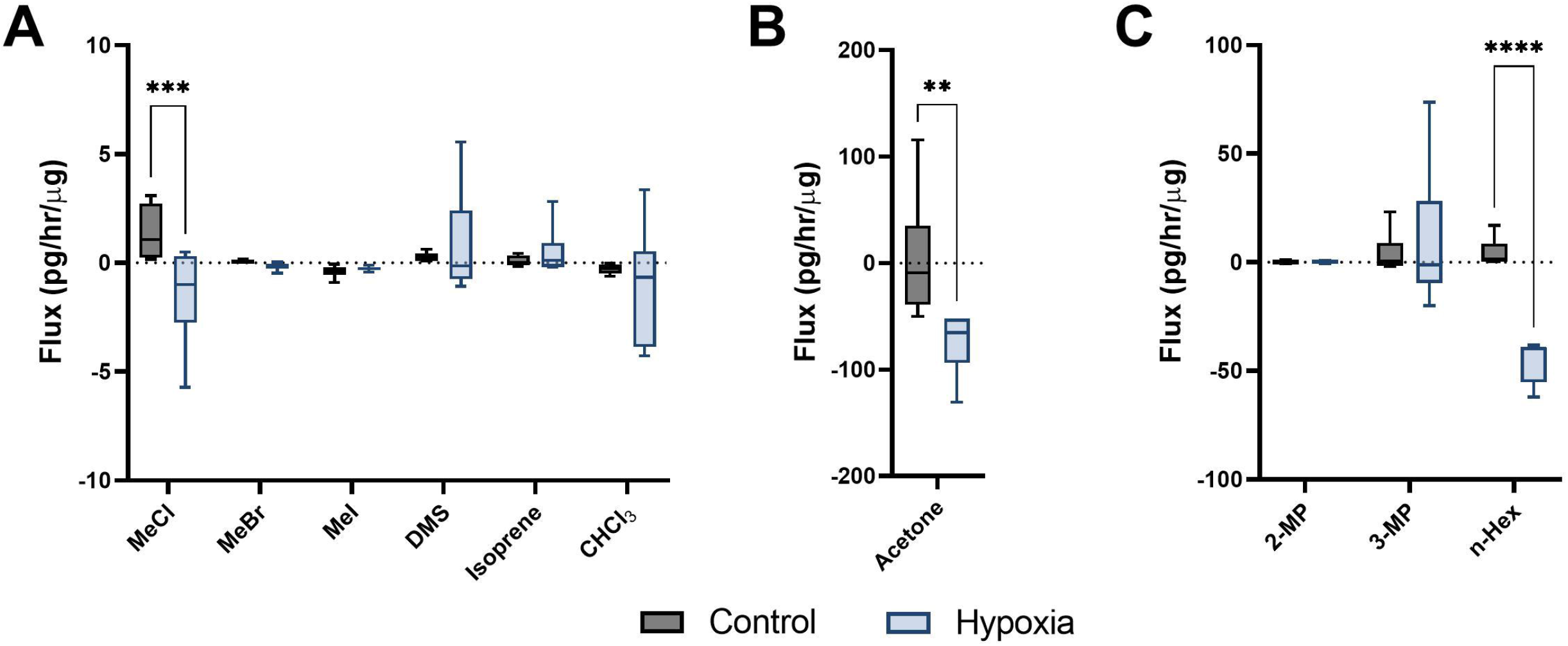
Cellular volatile response to hypoxia. Volatile flux (pg/hr/µg) for MDA-MB-231 cells in control conditions or hypoxia (24hr). Media subtracted and protein normalised VOC flux for MDA-MB-231 control cells (n = 6) and cells in hypoxia (n = 6). CHCl3 = Chloroform, OMS = Dimethyl sulfide, MeBr = Methyl bromide, MeCI = Methyl Chloride, Mel = methyl iodide, MeSH = Methanethiol, 2-MP = 2 methyl pentane, 3-MP = 3 methyl pentane, n-Hex = n-Hexane. Boxplot whiskers show median ± Tukey distribution, n = 6. ANOVA followed by Bonferroni post hoc test was performed; ** p < 0.01; ***p < 0.001; ****p < 0.0001.

### Production of Styrene under hypoxic conditions

Cells maintained under hypoxic conditions significantly produced styrene as determined by non-targeting GG-MS approaches (Figure 3A). Styrene was not found in the headspace of control cells (ND, or not detected) and styrene fluxes in media blanks were not significantly different from zero, while fluxes from hypoxic cells were significantly different from both hyperoxic controls and media blanks. Styrene was identified through spectral matching, followed by known standard injections. No other compounds were found to be significantly altered using the non-targeted SCAN method.

**Figure 3.**
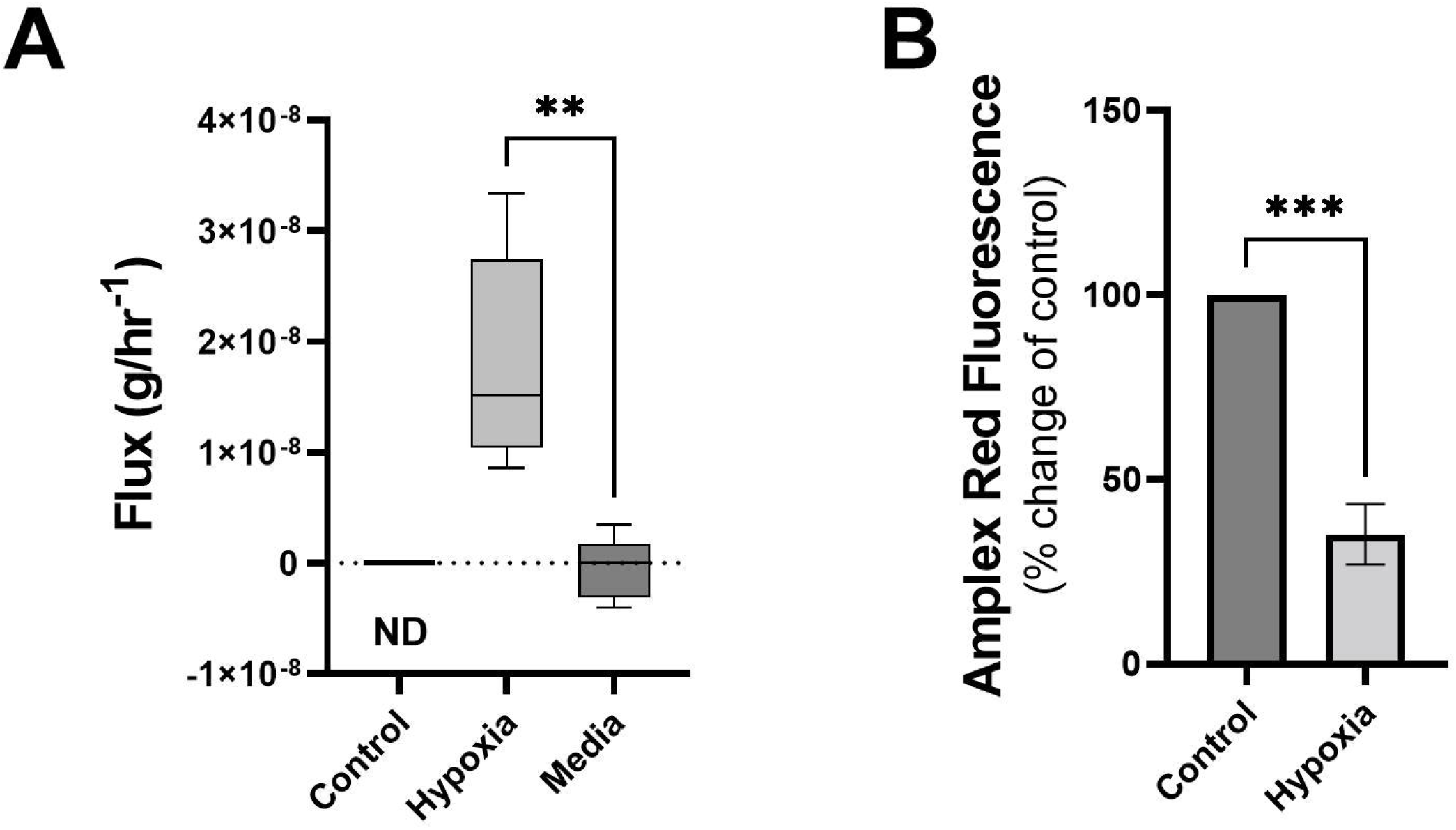
Cells under hypoxic conditions produce styrene and exhibit reduced ROS. **(A)** Volatile flux (g/hr^-1^) for styrene from MDA-MB-231 cells in control conditions or hypoxia and media only (24hr). Styrene was Non Detected (ND) for control cells. **(B)** Amplex Red assay was performed following 24 hours incubation as a measure of reactive oxygen species (ROS), H_2_O_2_. Shown as percentage change from relative control. Boxplot whiskers show median± Tukey distribution, A; n = 6. Student’s T-test was performed for A (hypoxia vs media) and B, ** p < 0.01, ***p < 0.001.

### Reactive oxygen species are reduced under hypoxia

Changes in volatiles, including alkanes, have been linked to increases in ROS (25). The observed uptake of n-Hexane in hypoxic MDA-MB-231 cells could therefore be correlated with alterations in ROS levels in these cells. Following 24 hours exposure to hypoxic conditions, ROS, as determined by Amplex Red assay, showed significant reduction compared to control (Figure 3B).

## Discussion

We demonstrate that the static headspace sampling chamber is capable of maintaining a low oxygen environment for >24 hours, as evidenced by chamber concentrations and cellular ROS response. Furthermore, we show that VOCs from cells maintained under low oxygen conditions can be sampled, and that these cells produce a significantly different volatile profile than either media blanks or identical cells exposed to hyperoxic conditions.

Three out of 10 compounds targeted by SIM revealed quantifiable, differential metabolic responses in cells exposed to hypoxic conditions (1% O_2_) relative to those maintained in normal laboratory conditions (21% O_2_, physiological hyperoxia). Our previous results quantified alterations in MDA-MD-231 cells for these volatiles after treatment with the chemotherapeutic agent Doxorubicin. When placed under cellular stress through Doxorubicin treatment only MeCl showed a similar stress response (enhanced uptake). In contrast, neither acetone nor hexane (or hexane isomers) were consumed or degraded significantly [4].

Over 24 hours of doxorubicin treatment has been shown to increase ROS (26) whereas the opposite has been shown in cells maintained in hypoxic conditions (27). We demonstrate here a significant reduction in ROS in MDA-MB-231 cells following 24hrs of hypoxia (Figure 3B). Cellular stress response mechanics and differences in cellular state could therefore be identified and quantified through volatile metabolic approaches. Alkanes have been positively correlated with ROS previously (25), and here we demonstrate a decrease in n-hexane within hypoxic cells (Figure 2C) with diminished ROS content while in cells treated with doxorubicin we observed non-significant increases (4).

Acetone has been associated with glucose metabolism and is a by-product of lipolysis in the breath (28, 29). Increased uptake of glucose has been shown under hypoxia (30) and decreases in acetone in the breath are observed following high glucose foods (29). Here, we have observed increased uptake of acetone under hypoxic conditions (Figure 2B).

The production of styrene by cells under hypoxia could be a defining VOC biomarker for cancer since hypoxia is characteristic of the tumour microenvironment (17). Our recent review showed that, despite substantial variability in reported outcomes, aromatics are powerful descriptors of cancer (3). Five studies have previously reported styrene in the breath of lung cancer patients using non-targeting approaches (31, 32, 33, 34, 35, 36). Styrene has also been reported as higher in the breath of lung cancer patients in studies using other approaches (37, 38, 39). However, styrene has been shown to be higher in the breath of smokers (34) and so is often considered, along with other aromatics compounds, to be a confounding contaminant since high percentages of lung cancer patients have a history of smoking. (3). Styrene has also been reported in the breath of patients with ovarian (40), gastric (41, 42) and liver (43) cancers.

Styrene utilisation as a breath-based diagnostic biomarker may be challenging since environmental contamination would need to be considered (11). Our outlined method accounts for environmental VOCs through a flux analysis that incorporates two temporal sampling points, a starting sample following equilibration with the local atmosphere and a second sample at a later time point. This allows us to determine when available environmental volatiles are being added to (metabolically produced) or consumed/degraded by cells. This is important where environmental VOCs may mask effects or differences, such as high traffic, urban environments or perfumed indoor spaces.

Environmental-correction sampling approaches such as this chamber headspace method may present an opportunity to overcome challenges to applications within the clinic, particularly with breath samples taken from ambient air as well as exhalate from the patient. The two time point sampling approach is particularly important since production of compounds with large initial concentrations, or consumption/degradation of compounds are often challenging to detect using single time point sampling methods.

We have observed that cellular consumption of VOCs (MeCl, Acetone and n-Hexane) is descriptive of hypoxic stress and that chemotherapeutic stress also induces consumption of VOCs (4); notably MeCl. To our knowledge this is the first example of a controlled environment experiment performed under low oxygen conditions that both a) quantifies VOC fluxes from a cellular model and b) utilises a VOC injection of gases to monitor ongoing anaerobic metabolism of compounds. We have demonstrated a novel method for induction and maintenance of low oxygen for the study of volatile fluxes. This approach allows new dynamics to be explored for the discovery of cell to patient translational biomarkers. It is perhaps worthy of note that many of the published methods for breath research would not have identified or quantified the methyl chloride or hexane results, due to the small changes (pptv) observed.

We have previously shown how ‘volatile metabolic flux’ can separate cell type and response to chemotherapeutic stress (4). This chamber-based method has also been successfully used with mice models, quantifying both mouse-breath and faecal volatiles (4). Here, we demonstrate how this chamber-based approach can identify cells under hypoxic stress. Using a novel method to identify hypoxia-induced VOCs, we demonstrate potential biomarkers of cancer. Importantly these biomarkers are both produced and consumed by cells under hypoxic stress. MeCl, acetone, n-hexane and styrene are clinically interesting compounds requiring further investigation. The compounds reported here have been reported as present in human breath (44) and we have shown that these compounds vary in response to cellular stress, from previously published doxorubicin (4) and here, hypoxic stress. Together this suggests they are able to differentiate cellular response due to pathophysiological differences. These compounds are from diverse functional chemical groups and we have previously demonstrated the ability of functional chemical groups to separate disease groups with greater ability than individually considered compounds (3). A functionally diverse group of VOCs could give greater power when building a ‘breath-print’ for diagnosis (3).

## Conflict of Interest

The authors declare that the research was conducted in the absence of any commercial or financial relationships that could be construed as a potential conflict of interest.

## Author Contributions

Conceptualization, T.I., W.J.B. and K.R.R.; Data curation, T.I.; Formal analysis, T.I.; Funding acquisition, S.T.S., W.J.B. and K.R.R.; Investigation, T.I.; Methodology, T.I., M.R. and K.R.R.; Project administration, T.I. and S.T.S.; Resources, M.R., W.J.B.; Visualization, T.I.; Writing— original draft, T.I.; Writing—review and editing, S.T.S., W.J.B., M.R. and K.R.R. All authors have read and agreed to the published version of the manuscript.

## Funding

This research was funded by the White Rose Mechanistic Biology Doctoral Training Program, supported by the Biotechnology and Biological Science Research Council (BBSRC) BB/M011151/1.

## Acknowledgements

The authors would like to acknowledge the support provided by Mark Bentley in the University of York Department of Biology workshop. This article was submitted as a preprint to BIORXIV prior to submission to.

## Notes

### Competing Interest Statement

The authors have declared no competing interest.

